# Computational Prediction of Disease Detection and Insect Identification using Xception model

**DOI:** 10.1101/2021.08.10.455608

**Authors:** Lidia Cleetus, A Raji Sukumar, N Hemalatha

**Affiliations:** St Aloysius College of Management and Information Technology; TECHTERN Pvt. Ltd; St Aloysius college of Institute of Management and Information Technology

**Keywords:** convolutional neural network (CNN), Deep learning, plant disease and pests, transfer learning.

## Abstract

In this paper, a detection tool has been built for the detection and identification of the diseases and pests found in the crops at its earliest stage. For this, various deep learning architectures were experimented to see which one of those would help in building a more accurate and an efficient detection model. The deep learning architectures used in this study were Convolutional Neural Network, VGG16, InceptionV3, and Xception. VGG16, InceptionV3, and Xception are categorized as the pre-trained models based on CNN architecture. They follow the concept of transfer learning. Transfer learning is a technique which makes use of the learnings of the models previously trained on a base data and applies it to the present dataset. This is an efficient technique which gives us rapid results and improved performance. Two plant datasets have been used here for disease and insects. The results of the algorithms were then compared. Most successful one has been the Xception model which obtained 82.89 for disease and 77.9 for pests.

## I. Introduction

Plant diseases and pests have established themselves as one of the significant challenge to the agricultural sector by becoming a major threat to food security. And due to lack of proper infrastructure, they are yet to be rapidly identified in various parts of the world. A faster and accurate detection of disease and pest in the plants can enable the development of an early treatment technique and at the same time reduce the economic loss faced by agriculturists. The plant damaging diseases and insect pests are not only a food security threat but they can also result in terrible repercussions for farmers whose daily bread depends on agriculture [2].

The agriculture sector has huge potential to improve the demand for food and supply nutritious and healthy food. For farmers, insect identification is a challenging task since a significant portion of the crops get damaged and the quality of the crops gets degraded as a result. The drawback with traditional insect identification is that it requires well trained taxonomists to accurately identify the insects based on their morphological features [3].

An agricultural pest can be described as an insect population which feeds from plant tissue resulting in commercial damage. Majority of the pests are mites or insects. The growth of a pest will depend majorly on the pressure on the external insect, the local weather, greenhouse blueprint and the crop management practices. A pest can be catastrophic resulting in losses across various fronts like fiscal (loss of income, profit and investments), social (depopulation in backward areas), and psychological (panic and commotion). The existence of pests and diseases does pose threat for the farmer owning the farm but it also poses a threat for the nearby and at times distant holdings. Rapid pest identification is of utmost importance in terms of yield productivity as well as in reducing the pesticide usage. Eye observation methods by human beings have been in usage for many years but they are not productive when it comes to large crops [4].

Deep Convolutional Neural Network has been made use for detection of disease and pest identification. Large neural networks can be time consuming when put in training but once trained we can quickly classify images. This attribute makes them more suitable for smartphone consumer applications. While training neural networks, focus will be on improving the mapping during the whole process which is done by tuning certain parameters in the network. This process is computationally challenging. In recent times, both conceptual and engineering breakthroughs have been used to dramatically improve this process.

Conventional image recognition and classification methods of mechanical design features are capable of only extracting the underlying features. It is a hard task to extract the complex and deep image attribute information. Deep learning methods can solve this problem. To acquire multi-level image attribute feature information such as high-level, intermediate and low-level features, the deep learning techniques will perform unsupervised learning directly on it. Conventional disease and pests detection algorithms largely adopt the image recognition methods of manually designed attributes which is difficult as well as it depends on luck and experience. It is also unable to learn and extract features from the original image by itself. Performances of these kind of approaches depends on the pre-defined attributes. Feature engineering is a difficult process that needs to be approached again in case the problem and the dataset changes. On the contrary, deep learning has the ability of learning the features automatically from huge data without any manual intervention. The deep learning model comprises of many layers which has feature expression caliber and good autonomous learning ability, and can by itself perform feature extraction on original images. Therefore, deep learning has the prospective to play an important role in the field of image recognition of diseases and pests [5].

This paper has been distributed into four sections. Section two talks about the materials used and methodology followed in the paper. Results are discussed in section three and paper is brought to conclusion in section four.

## II. Materials and Methods

This section covers the materials used in this study. It also explains the work done in this study in more detail.

### A. Data Source

The data has been taken from the Kaggle repository. We have used two datasets in our study – Tomato Leaf Disease and Pests Identification. Both of the datasets consists of images of different species of diseases in tomato and pests respectively. There are 10,000 images in the disease dataset while the pest dataset consists of 1,669 images (Figure 1 and 2).

**Fig. 1.**
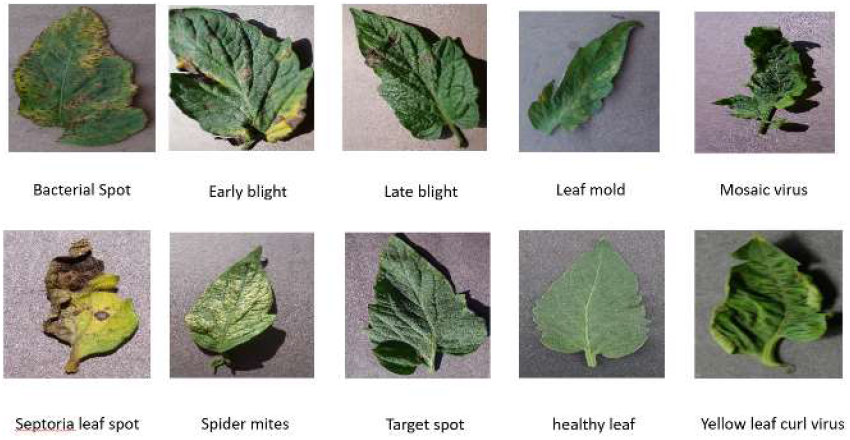
Sample images of Tomato Leaf Disease dataset

**Fig. 2.**
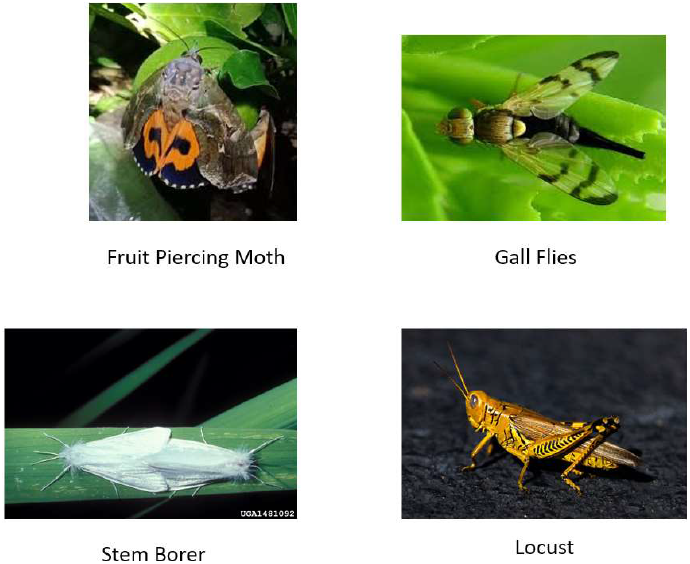
Sample images of Pest dataset

### B. Features

The Tomato leaf disease dataset contains 10 classes including 9 different diseases found in tomato leaves and 1 healthy leaf class. The diseased classes include Bacterial Spot, Mosaic virus, Early blight, Late blight, Septoria leaf spot, Leaf mold, Spider mites, Yellow leaf curl virus, and Target spot. The Pests identification dataset contains 4 classes *viz*., Fruit Piercing Moth, Gall Flies, Locust, and Stem Borer. Sample images of each class can be seen in Figure 1 and 2.

### C. Data Preprocessing

Data pre-processing is a data mining technique that transforms raw data into an understandable format. Typically, a real world data is always inconsistent, incomplete, and inaccurate (contains outliers or errors). Such kind of data cannot be sent through a model. That would cause certain errors. That is why we need to preprocess data before sending through a model. There are several steps we need to follow for data preprocessing [6]:

#### 1) Data acquisition

We must first acquire the relevant dataset to build and develop the deep learning models. The dataset comprises of data collected from various multiple sources. It is then integrated into a proper format to construct a dataset. Formats of dataset differs according to the use cases. For example, a medical dataset will differ completely from a business dataset. Medical dataset will include healthcare-related data whereas a business dataset comprises of relevant business and industry related data. Here, we have acquired image datasets for the purpose of detecting and identifying the diseased leaves and pests.

#### 2) Import the relevant libraries

Python has established itself to be the most preferred and extensively used library around the world by Data Scientists. The predefined Python libraries are majorly used to perform certain data preprocessing jobs. The main libraries used in Deep Leaning data preprocessing is:

- Numpy: Python’s basic scientific computation package so you can put any kind of math operation in your code. NumPy also allows us to add huge multi-dimensional arrays and arrays in code.
- Pandas: An open-source library for manipulating and analyzing data in Python. Pandas is widely used to import and manage datasets. Has convenient, powerful data structures and data analytics tools making it one of the powerful data analysis library.
- Matplotlib: A 2D Python plot library for plotting different types of charts in Python. It offers some of the finest charts in diverse print formats and is known for its interactive environments on all platforms.
- Scikit-Learn: Scikit-learn (formerly known as scikits.learn and also as sklearn) is a ML library for the Python language available for free. Contains numerous classification, regression, and clustering algorithms including Random Forests, K-means, Gradient Boosting, Support Vector Machines, and DBSCAN and is capable of working together with Python’s NumPy and SciPy scientific and numeric libraries [7].
- TensorFlow: TensorFlow is a free, open source software library. It can be used for a variety of tasks, but has a special focus on deep neural network training and inference. Tensorflow is a symbolic math library based on data flow and differentiable programming. Used for both research and production in Google, TensorFlow was developed by their Brain team for Google’s internal use.
- Keras: Keras is an open source software library. Keras provides a Python interface for ANNs. It poses as an interface to the TensorFlow library. It is designed to allow for quick experimentation with deep neural networks and focuses on being easy to use, scalable, and extensible. As part of the research efforts of the ONEIROS (Open NeuroElectronic Intelligent Robot Operating System) project, Keras was developed and its lead author is François Chollet, a Google engineer. Chollet is also the one who introduced the Xception model.
- OpenCV: A cross-platform library that is used to develop real-time CV (computer vision) applications. Its main focus is on image processing, video recording and analysis, including functions such as facial recognition and object recognition [8].

#### 3) Read the data

Using OpenCV/Matplotlib/PIL, we can easily read the images and plot it accordingly.

#### 4) Label Binarizer

A SciKit Learn class that takes categorical data as input and returns a numpy array. In contrast to Label Encoder, it encodes the data in dummy variables that indicate the presence of a certain label or not. Here we apply the LabelBinarizer() to the class labels [9].

#### 5) Conversion to numpy array

Images are converted into Numpy Array in Height, Width, Channel format so that further manipulation can be done on them.

#### 6) Rescale

Images are scaled down by factor 255 before feeding to the model. Some images have a high pixel range and some have a low pixel range. The images are all sharing the same model, weights and learning rate. There will be stronger loss in a high range image as compared to the weak loss found in a low range image. Their sum will contribute to the back propagation update. When all images are scaled to the same range [0,1], the images contribute more evenly to the total loss.

#### 7) Data Augmentation

An approach that creates modified versions of images in the dataset in order to artificially increase the size of a training data. This approach will apply various transformations to create different versions of images that belongs to the class same as the original image. Transformations encompass a number of image manipulation operations such as shifting, mirroring, zooming, and much more. In the real-world scenario, there might be a collection of images that were captured under limited conditions. The target application can exist in various conditions like a different position, brightness, orientation, scale, etc. These situations can be explained by training the neural network with the addition of synthetically modified data. *[10].*

## D. Data Splitting

The train_test_split function from the Python package Sklearn model selection can be used to split the data into train and test datasets. The split ratio of 70:30 is used for this study.

## E. Algorithms

### 1) Convolutional Neural Network (CNN)

This deep learning architecture consists of multiple layers or blocks – Convolutional Layer, Pooling Layer, Activation Function, Fully Connecter Layer, Loss Functions (Figure 3) [11].

- Convolution Layer - In this architecture, the most important part is the convolution layer, which comprises of a set of convolution filters (or kernels). The image passed as the input which is expressed as N-dimensional metrics, is convoluted with this set of filters to produce the output characteristics map.
- Pooling Layer – Subsampling of the feature maps is the major task of a pooling layer. These maps are produced after the convolution operations. To put it simply, this approach will reduce and create smaller feature maps from larger feature maps. It maintains the significant information (or properties) at each pace of the pooling level. Akin to the convolution operation, both the kernel and the stride are originally sized before the operation pooling is performed. There exist different types of pooling methods like Tree Pooling, Closed Pooling, Average Pooling, Minimum Pooling, Maximum Pooling, Global Maximum Pooling, and Global Average Pooling (GAP). The best known and most widely used pooling methods are maximum, minimum and GAP pooling (Figure 4).
- Activation function – In a neural network, the central function of all types of activation functions is to map the input to the output. The input value can be decided by calculating the weighted sum of the inputs along with the bias. The activation function has to make a decision concerning whether a neuron should be fired with note to a certain input or not by generating the corresponding output. Once all the weighted layers in CNN architecture are used, the nonlinear activation layers are deployed. The activation layers’ non-linear performance implies that the input-to-output mapping is of non-linear nature; In addition, CNN is given the opportunity to learn specific complex things by the layers. The activation function must be capable of differentiation, an exceedingly important feature since it will allow back propagation to be used in training of the network.
- Fully Connected Layer - This layer can be found at the end of a CNN architecture. In a fully connected layer, each neuron connects to all neurons present in the previous layer. It is used as a CNN classifier. Follows the fundamental method of the convolutional MLP neural network as it is a kind of feedforward ANN. The input will come from the last pooling or convolutional layer. Input in the vector form is generated using feature maps after it is flattened. The output from the FC layer constitutes the final output from CNN, as shown in Figure 3.
- Loss functions - The output layer which is the last layer in a CNN architecture performs the final classification. To compute the predicted error generated from the training samples, the output layer utilizes a few loss functions. The error is basically the difference between an actual and a predicted output. Through the CNN learning process, this error is optimized. Even so, the loss function will use two parameters to calculate the error. The first parameter of CNN is the estimated output also called as prediction. The second parameter is the actual output also called label.

**Fig. 3.**
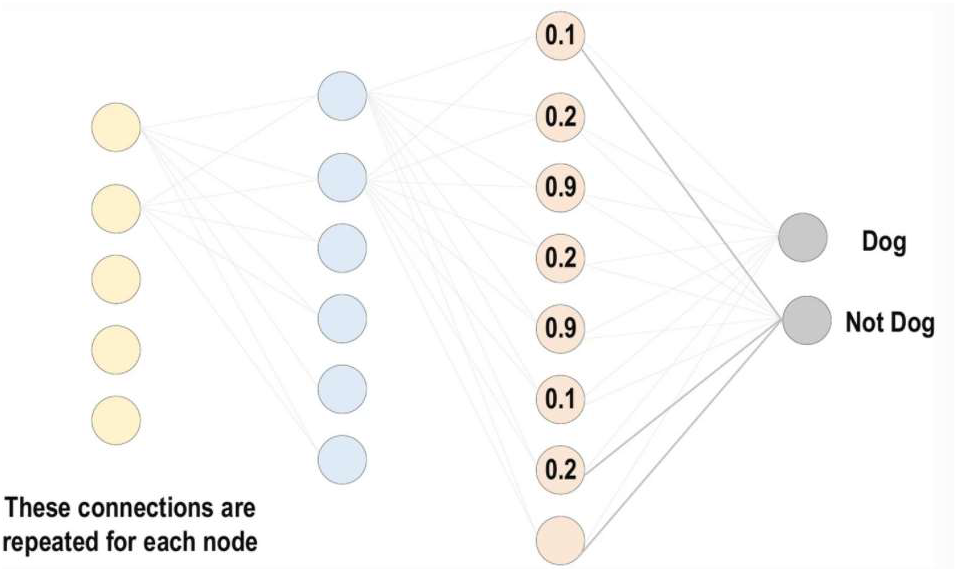
Architecture of CNN

**Fig. 4.**
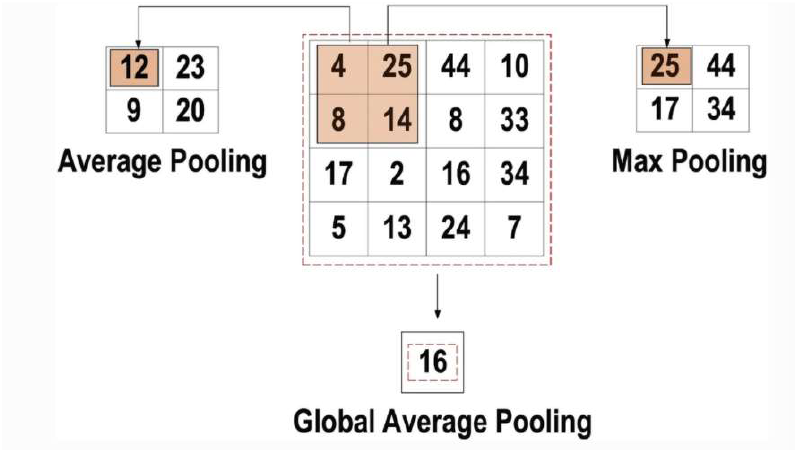
Different types of pooling layers

## F. Models

### 1) VGGNet

In their paper “Very deep convolutional networks for large-scale image recognition”, A.Zisserman and K.Simonyan of Oxford University proposed a CNN model – the VGG16. The model achieved 92.7% of accuracy in the Top5 tests by ImageNet, a dataset consisting of 1000 classes and more than 14 million images. VGG16 became one of the most noted models presented at ILSVRC 2014. It improves over the AlexNet model by substituting larger filters (11 in the first convolutional layer and 5 in the second one) with numerous 3×3 kernel size filters one after the other. NVIDIA Titan Black GPUs were used in the training of VGG16 for weeks (Figure 5) [12].

**Fig. 5.**
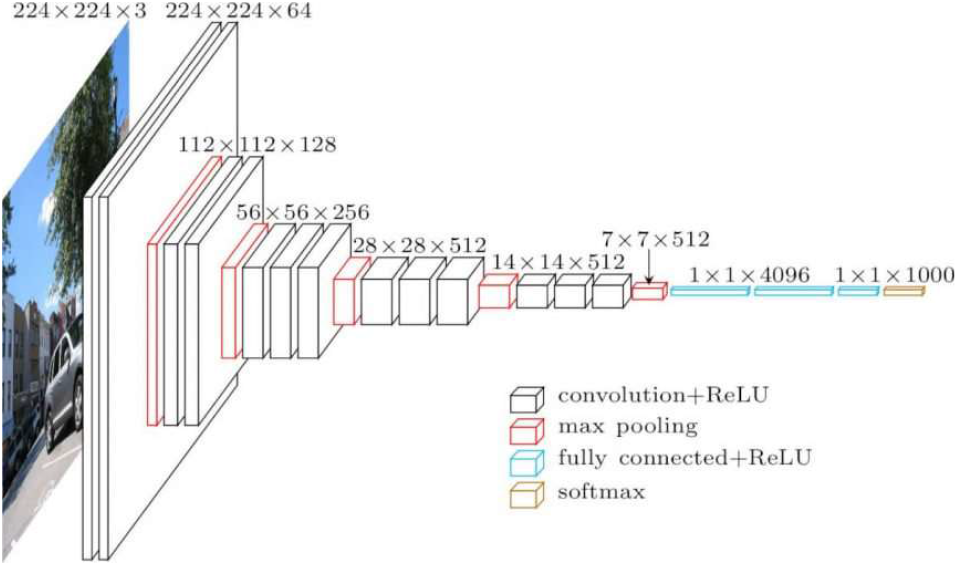
Architecture of VGG16

An RGB image of fixed size 224 × 224 is the input for the conv1 layer. The input passes through a stack of layers, using the filters having very small receiving field of 3 × 3. It also uses 1 × 1 convolution filter in one of its settings. This can be seen as a linear transformation of the input channels. The stride in the layers is fixed at 1 pixel. After convolution, the spatial resolution is retained due to the spatial padding of convolutional layer input. 5 layers of max pooling performs the spatial pooling, which corresponds to some of the convolutional layers. Max pooling is performed with a stride of 2 in a 2 × 2 pixel window.

An assembly of convolutional layers is followed by a three fully connected layers. First two of the fully connected layers have 4096 channels each whereas the third connected layer will have 1000 channels – one for each class and performs the 1000 way ILSVRC classification. Last layer comprises of the Softmax layer. All networks have the same configuration for the fully connected layers.

ReLU (rectification non linearity) is present in all hidden layers. Networks do not contain a LRN (local response normalization). These kind of normalization won’t improve the ILSVRC data performance, although it leads to higher memory consumption and computing time.

The configurations of ConvNet are described in detail below in Fig. 6. The names (A-E) refer to the nets. All configs keep to the architecture’s generic design and only differ in depth: from the A network comprising of 11 weight layers (8 conv and 3 FC) to the E network comprising of 19 weight layers (16 conv and 3 FC). The width of the layers (the number of channels) is quite small, beginning with 64 in the starting layer expanding till it reaches 512. After each max pool layer, the increase transpires by a factor of 2.

**Fig. 6.**
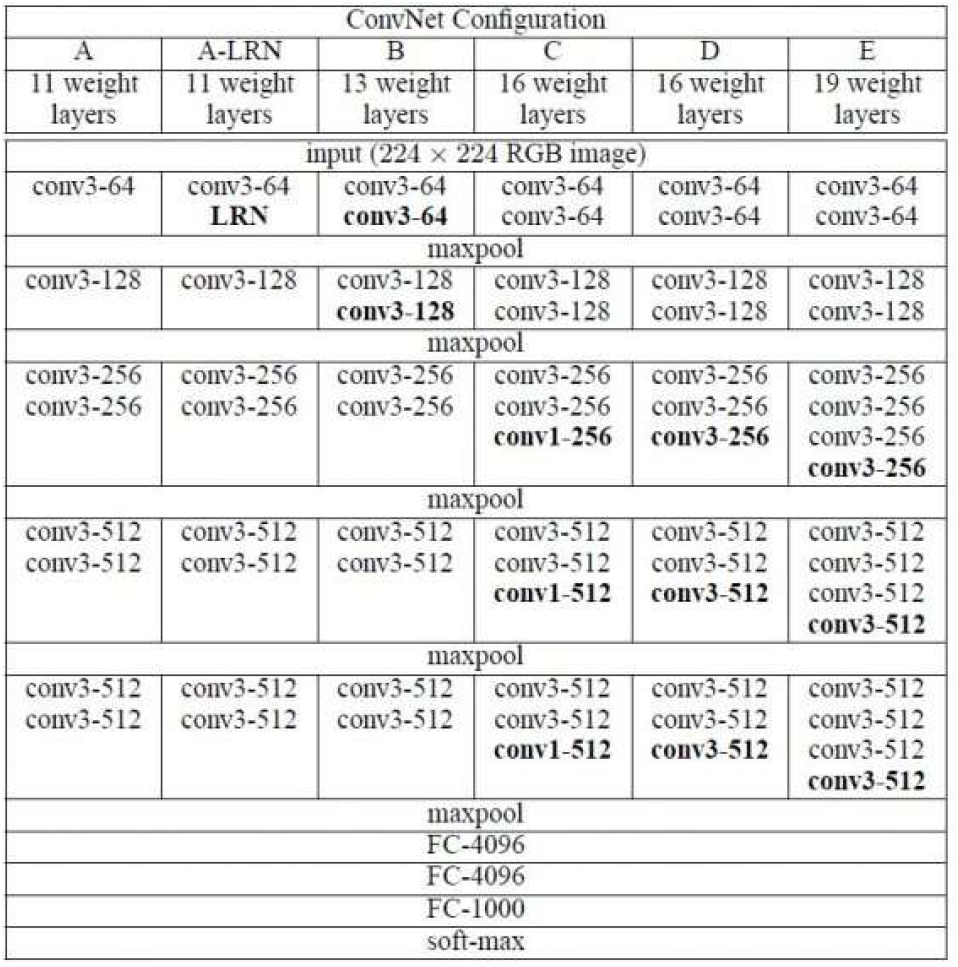
ConvNet configurations

### 2) InceptionV3

Inception v3 is primarily focused on consuming less computing power by modifying previous Inception architectures. It was in the article ‘Rethinking the Inception Architecture for Computer Vision’ where the idea made its first appearance. This article was published in the year 2015 and co-authored by V. Vanhoucke, S. Ioffe, J. Schens and C. Szegedy.

Inception Networks like GoogleNet or Inception v1 have been established to be more computationally efficient compared to VGGNet, both in terms of count of parameters generated and the economic costs involved. In an Inception network, it must be ensured that the computational advantages are not lost. Due to the insecurity of the new network, problem arises when an Inception network is being adapted for different use cases. In an Inception v3 model, various techniques were proposed to optimize the network in order to relax constraints and to facilitate model adaptation. Techniques include factorized convolution, regularization, dimension reduction, and parallel computations.

[13] Inception V3 is similar to and contains all the functions of the predecessor Inception V2 with following changes/additions:

- RMSprop optimizer was used.
- Batch Normalization in the FC layer of Auxiliary classifier.
- Usage of 7×7 factorized Convolution
- Label Smoothing Regularization: It is a method of regularizing the classifier by estimating the effect of skipping labels during training. Prevents the classifier from predicting a class with too great a degree of certainty. Adding label smoothing provides a 0.2% improvement over the error rate.

An Inception v3 network is built gradually, as explained below [14]:

- Factorized Convolutions: reduces the number of parameters participating. As a result, the computational efficiency and the network efficiency are optimized.
- Smaller convolutions: smaller convolutions replace larger ones resulting in quick training. Say, two 3 × 3 filters will replace a 5 × 5 convolution having 25 parameters. The resulting convolution will only have 18 (3*3 + 3*3).
- Asymmetric convolutions: A 1 × 4 convolution can replace a 4 × 4 convolution followed by a 4 × 1 convolution. Now, if a 3 × 3 convolution was replaced by 2 × 2, then the no of parameters will be slightly larger than the proposed asymmetric convolution.
- Auxiliary classifier: During training, an auxiliary classifier is inserted between layers. The resulting loss is then added to the loss of main network. In Inception v3, an auxiliary classifier will behave as a regularizer whereas in GoogLeNet, they were utilized for a deeper network.
- Grid size reduction: Usually the pooling operations reduce the size of the grid. However, to address computational cost bottlenecks, a more efficient technique is suggested, as shown in Fig. 8.

**Fig. 7.**
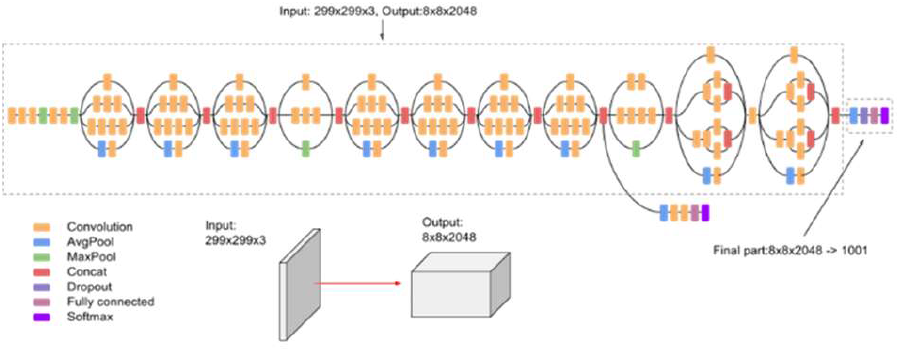
Architecture of Inceptionv3

**Fig. 8.**
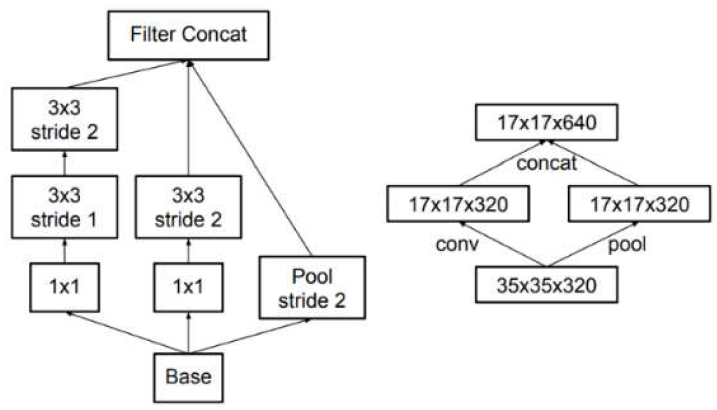
Grid size reduction

### 3) Xception

Xception, the extreme version of Inception, is an architecture based on depth-wise separable convolutional layers. With this modified depth-wise separable convolution it is even better than Inceptionv3 [15].

Xception was suggested by none other than François Chollet himself, the creator and administrator of the Keras library. The standard Inception modules are replaced by the Xception architecture which is an extension of Inception architecture. It’s a simple and modular architecture.

The architecture has 36 convolution layers that form the basis for extracting functions from the network. These 36 layers are structured into 14 modules, all of which with the exception of the first and last module have residual linear connections around them. The depth wise separable convolution layers will make up a linear stack with residual connections. As a result, it is easier to define and modify the architecture. When using a high-level libraries like Keras or TensorFlow Slim, only 30 to 40 lines of code are required, similar to an architecture like VGG16, but in contrast to architectures like Inception V2 or Inception V3, which are much more complex to define (Figure 9).

**Fig. 9.**
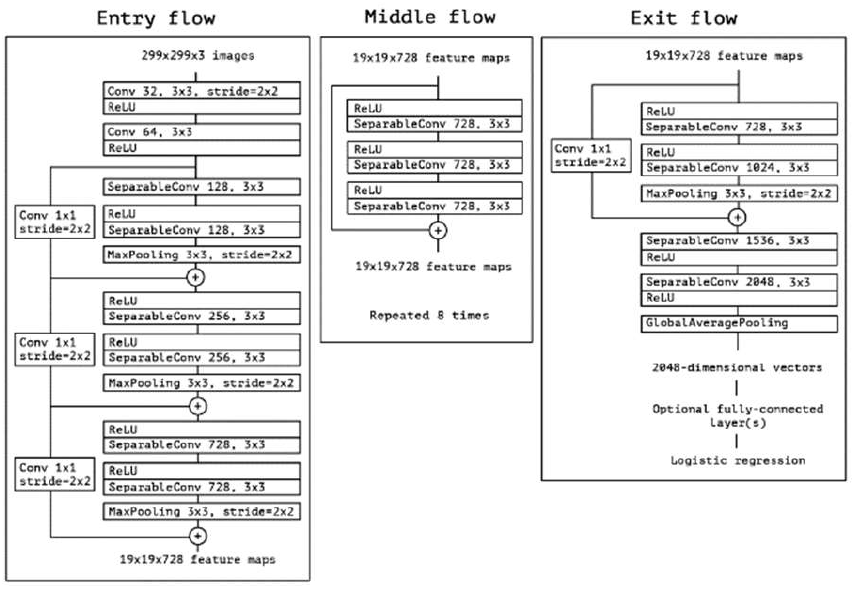
Architecture of Xception

As in the Fig. 9, the modified deeply separable convolution seen is SeparableConv. These SeparableConvs are placed in the deep learning architecture and are regarded as inception modules. And there are leftover connections (or shortcuts / skips) placed for all flows originally suggested by ResNet. There are residual connections as you can see from the architecture. Here, it uses the non-residual version to inspects for the Xception.

The modified separable convolution is the pointwise convolution which is followed by a depth wise convolution. The inception module inspired this modification such that the 1 × 1 convolution is performed before each spatial n × n convolution. There are slight differences between the original and the modified version which are listed below:

- The order of operations: The original depth-wise separable convolutions will first perform channel wise spatial convolution and then a 1 × 1 convolution. Whereas, the modified version will perform in the reverse order i.e. 1 × 1 convolution followed by channel wise spatial convolution. It becomes insignificant when used in stacked setting as only small differences will be visible in the beginning and end of all chained inception modules.
- The Presence/Absence of Non-Linearity: After the first operation, the original Inception module is free from linearity. And in the modified version of Inception, there is no inter-ReLU non-linearity.

## III. Results

For the Tomato leaf disease dataset, Xception model performed better amongst all model/architecture. It produced an accuracy score of 82.89%. CNN architecture scored an accuracy score of 82.83% making it the second best model. For the rest of the models, i.e. VGG16 and Inceptionv3, the test accuracy score produced was 82.11% and 80.33% respectively (Table 1). Accuracy and loss plots were plotted using the plot function from the matplotlib.pyplot package as shown in Fig. 11.

**Table 1:** Results table for disease dataset

| Architecture/Model | Test Accuracy (%) |
| --- | --- |
| CNN | 82.83 |
| VGG16 | 82.11 |
| INCEPTION V3 | 80.33 |
| XCEPTION | 82.89 |

**Fig. 10.**
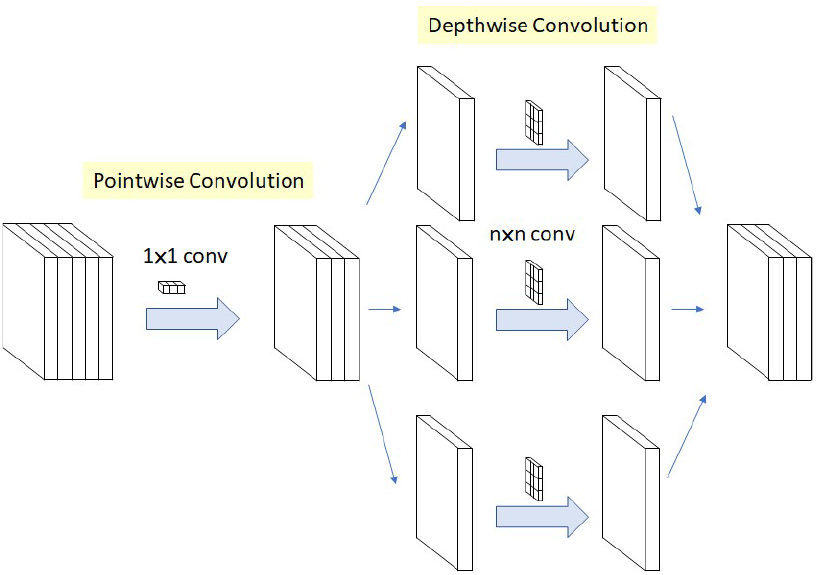
Modified depthwise separable convolution in Xception

**Fig. 11.**
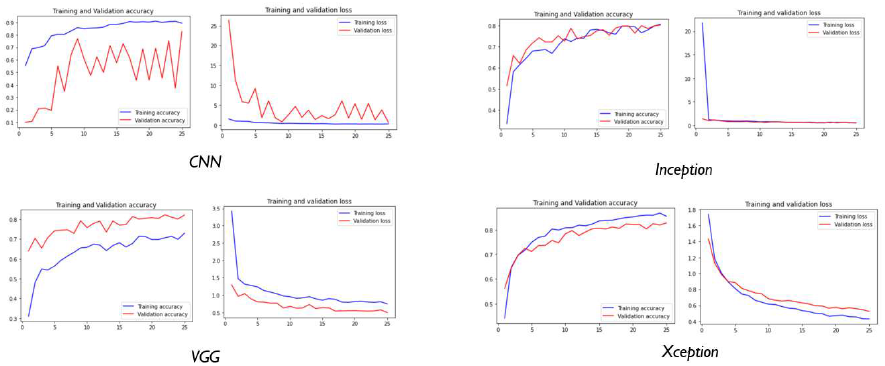
Accuracy and Loss plots for disease dataset

And for the Pest dataset, Xception model performed better amongst all model/architecture. It produced an accuracy score of 77.90%. Inceptionv3 model scored an accuracy score of 77.19% making it the second best model. For the rest of the networks, i.e. VGG16 and CNN, the test accuracy score produced was 71.74% and 24.28% respectively (Table 2). Accuracy and loss plots were plotted using the plot function from the matplotlib.pyplot package as shown in Fig. 12.

**Table 2:**
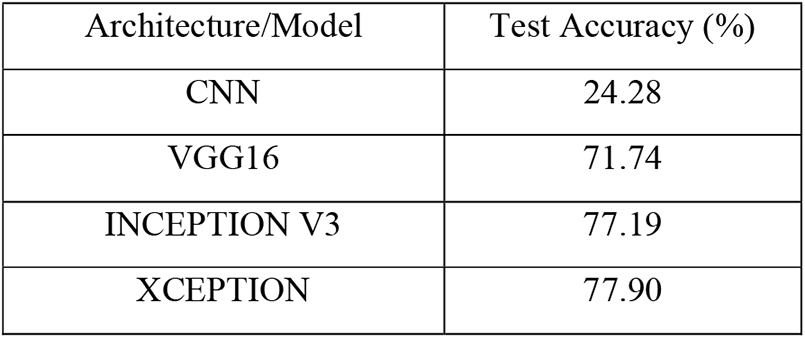
Results table for pest dataset

**Fig. 12.**
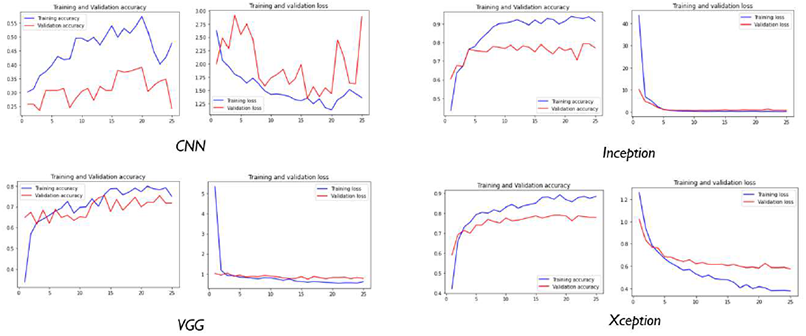
Accuracy and Loss plots for pest dataset

## IV. Conclusion

After comparing the performance of each model with regards to test accuracy score, it can be concluded that the Xception model will be the best choice for the detection and identification of tomato leaf diseases and pests. The accuracy achieved by the model is 82.89% for tomato disease dataset and 77.90% for the pest dataset.

## V. ACKNOWLEDGEMENT

The authors extend their appreciation to the Deputyship of RESEARCH AND INNOVATION wing of TECHTERN Pvt. Ltd. through SMART-AGRO research and for providing all support for this research work with the project number TTRD-DS-01-2021. This is also an extension of a post-doctoral research program under Kannur University.

## REFERENCES

[1] D. Gupta, “Transfer learning and the art of using Pre-trained Models in Deep Learning,” Analytics Vidhya, 01 June 2017. [Online]. Available: https://www.analyticsvidhya.com/blog/2017/06/transfer-learning-the-art-of-fine-tuning-a-pre-trained-model/.

[2] S. P. Mohanty, D. P. Hughes and M. Salathe, “Using Deep Learning for Image-Based Plant Disease Detection,” Frontiers in Plant Science, vol. 7, p. 1419, 2016.

[3] “Pest insects,” Agriculture and Food: Department of Primary Industries and Regional Development, [Online]. Available: https://www.agric.wa.gov.au/pests-weeds-diseases/pests/pest-insects.

[4] A. Ansuategi, L. Susperregi, C. Tubío, I. Rankić and L. Lenža, “A Benchmarking of Learning Strategies for Pest Detection and Identification on Tomato Plants for Autonomous Scouting Robots Using Internal Databases,” Journal of Sensors, p. 15, 2019.

[5] J. Liu and X. Wang, “Plant diseases and pests detection based on deep learning: a review,” 2021

[6] K. Goyal, “Data Preprocessing in Machine Learning: 7 Easy Steps To Follow,” 22 January 2020. [Online]. Available: https://www.upgrad.com/blog/data-preprocessing-in-machine-learning/.

[7] “Scikit-learn Tutorial: Machine Learning in Python,” Dataquest, 15 November 2018. [Online]. Available: https://www.dataquest.io/blog/sci-kit-learn-tutorial/

[8] “About OpenCV,” OpenCV, [Online]. Available: https://opencv.org/about/.

[9] M. A. Reddy, “Encoding Categorical data in Machine Learning,” Medium, 23 June 2019

[10] J. Brownlee, “How to Configure Image Data Augmentation in Keras,” Machine Learning Mastery, 05 July 2019

[11] L. Alzubaidi and J. Zhang, “Review of deep learning: concepts, CNN architectures, challenges, applications, future directions,” Journal of Big Data, 2021

[12] P. Jay, “Image Classification Architectures review,” Medium, 19 July 2018

[13] “Inception V2 and V3 – Inception Network Versions,” GeeksforGeeks, 16 July 2020

[14] V. Kurama, “A Review of Popular Deep Learning Architectures: ResNet, InceptionV3, and SqueezeNet,” PaperspaceBlog, 2020

[15] S.-H. Tsang, “Review: Xception — With Depthwise Separable Convolution, Better Than Inception-v3 (Image Classification),” Towards Data Science, 25 September 2018

